# Multi-omics analyses from a single sample: Prior metabolite extraction does not alter the 16S rRNA-based characterization of prokaryotic community in a diversity of sample types

**DOI:** 10.1101/2023.07.18.549538

**Authors:** Sébastien Duperron, Pierre Foucault, Charlotte Duval, Midoli Goto, Alison Gallet, Simon Colas, Benjamin Marie

## Abstract

Massive sequencing of the 16S rRNA gene has become a standard first step to describe and compare microbial communities from various samples. Parallel analysis of high numbers of samples makes it relevant to the statistical testing of the influence of natural or experimental factors and variables. However, these descriptions fail to document changes in community or ecosystem functioning. Non-targeted metabolomics are a suitable tool to bridge this gap, yet extractions protocols are different. In this study, prokaryotic community compositions are documented by 16S rRNA sequencing after direct DNA extraction, or after metabolites extraction followed by DNA extraction. Results obtained using the V3-V4 region on non-axenic cultures of cyanobacteria, lake water column, biofilm, gut of wild and lab-reared fish, indicate that prior extraction of metabolites does not influence the obtained image of prokaryotic communities. This validates sequential extraction of metabolites followed by DNA as a way to combine 16S rRNA sequencing with metabolome characterization from a single sample. This approach has the potential to complement community structure characterization with a proxy of their functioning, without the uncertainties associated with the use of separate samples.

## Introduction

Omics approaches allow analyzing genomes, transcripts, lipids, proteins, and metabolites to an unprecedented level of depth. Each is informing cellular processes and molecular responses at particular time and size scales, making them highly complementary. The next big step forward is thus to combine them into multi-omics approaches. Correlating multi-layered datasets comes with major technical, statistical and conceptual challenges (Krassowski et al., 2020; Tarazona et al., 2021). One basic challenge not to be neglected, particularly relevant to microbiologists, is the fact that these approaches are often destructive, and thus cannot be routinely performed from a single small sample and require parallel extraction on split samples (*e*.*g*.(Clerissi et al., 2023)).

One of the most commonly applied approaches in microbial ecology is the characterization of microbial community compositions using high-throughput sequencing of ribosomal RNA-encoding genes from extracted DNA (Alivisatos et al., 2015; Dubilier et al., 2015). Despite it is widely used and highly informative, this approach providing a description of « who is there » is often criticized for yielding limited information about « what is going on » (Louca et al., 2018). Quick and easy ways to address the functioning of microbial systems with a desirable level of replication are wanting. In this context, non-targeted metabolomics has emerged as a fast, convenient and sensitive approach for fingerprinting the functioning of various biological systems (Colas et al., 2020; Le Manach et al., 2019; Sogin et al., 2019; Zhang et al., 2016). Indeed, metabolome composition of an organism or community can be used as a proxy of its phenotype, as it is a function of organisms (or holobionts) genotype and its interaction with biotic and abiotic factors. Metabolome composition can vary quickly, within seconds, in response to stimuli, stress, changes in the environment or overall health status including age, sex, disease and treatments (Khemtong et al., 2015; Rauckhorst et al., 2022). Metabolomics datasets are usually difficult to interpret on their own, in particular owing to the lack of proper databases that could link peaks identified in mass spectrometry to particular metabolites, even more so in the microbial world (Gertsman and Barshop, 2018). However, these are particularly relevant when combined in multi-omics approaches, for they can provide a quick measure of functional variations that can be linked to another omics, as for example recently performed for transcriptome analysis (Madaj et al., 2023).

Once metabolites are extracted from a sample, for example with classical methanol extraction procedure (Prasad Maharjan and Ferenci, 2003), other types of molecules can be recovered from the remaining pellet including proteins, lipid membranes and DNA (Roume et al., 2013; Valledor et al., 2014). However, the effect of prior methanol extraction of metabolites on the datasets obtained when analyzing these other molecules has rarely been investigated so far. Interestingly, a recent study has shown that RNAseq dataset quality was only marginally affected by prior metabolite extraction with 80% methanol in mice model, suggesting that single sample combination of metabolomics and transcriptomics was tractable in this case (Madaj et al., 2023).

For microbiologists, being able to produce various omics data from a single sample is particularly relevant. Indeed, microbial habitats are heterogeneous at a micrometer scale, for example in the multiple layers of a biofilm, in micro niches in soils, or in different regions of a host’s gut (Nunan, 2017; Shaffer et al., 2022). Splitting samples prior to applying different omics approaches to distinct subsamples thus potentially hampers our ability to apply correlation algorithms on results from the obtained datasets if samples are heterogenous. Single sample characterization of metabolites and prokaryotic community compositions could offer a quick and efficient way to correlate changes observed in the composition of communities and metabolome, linking community structure to functional fingerprinting. Besides this evident complementarity, both methods are time- and cost-effective, allowing comparison of numerous samples, and thus replication levels that allow proper hypothesis testing (Singh et al., 2019). Various recent studies have emphasized the potential of combining these approaches, one remarkable example identifying taxa and metabolites specific to different environments in a large scale study of earth microbiome (Shaffer et al., 2022). Although most of these studies did split samples in aliquots dedicated to each of the methods (Nothias et al., 2023; Shaffer et al., 2022), a few have performed combined extractions of metabolites and DNA including by ourselves (Foucault et al., 2022; Gallet et al., 2023). However, the absence of potential bias introduced by this method using internal control still remains to be demonstrated.

In order to investigate whether it is possible to use a single sample to perform both metabolomic and prokaryotic community analyses without affecting results from the latter, the aim of this study is to test whether prior metabolite extraction modifies the recovered image of community compositions. For this, we analyzed community compositions using a standard 16S rRNA high-throughput sequencing approach (V3-V4 region) on two types of extractions: classical DNA extraction using a kit, or standard methanol metabolite extraction followed by the same classical DNA extraction performed on the pellet. Communities structure and compositions were compared after the two types of extraction on different types of samples that are commonly analyzed, some coming from lab settings (non-axenic cyanobacterial strains cultures and gut of lab-reared fish) and some from the environment (lake water, river biofilm, and gut of a wild fish). To our knowledge, this study is the first attempt to test whether single sample metabolite followed by DNA extraction introduces a bias in the obtained image of prokaryotic communities.

## Material and methods

### Sample preparation

Four clonal non-axenic cyanobacterial strains of *Microcystis aeruginosa* were obtained from the culture collection of cyanobacteria at the Museum national d’Histoire naturelle (Hamlaoui et al., 2022). Strains PMC-728.11, PMC-807.12, PMC-810.12 and PMC-826.12 were cultured in BG-11 medium at 25°C in 250-mL Erlenmeyer vessels, with a photon flux density of 12 mmol·m^2^·s^-1^ and a 12:12-h light/dark cycle and was lyophilized (24h at -50°C). Gut of six medaka fish (*Oryzias latipes*) from the Museum national d’Histoire naturelle rearing facility, and from one wild specimen of *Perca fluviatilis* from the Jablines lake (48°55 03.3 N 2°43 31.3 E) were dissected, flash-frozen in liquid nitrogen, pooled and ground to power using mortar and pestle, and lyophilized (24h at -50°C). Three distinct water sample were produced by filtering 500 mL of water from three lakes near Paris, namely Champs-sur-Marne (filter 1, 48°51 47.6 N 2°35 51.6 E), Cergy (filter 2, 49°01 29.5 N 2°02 47.0 E) and Jablines (filter 3, 48°54 48.3 N 2°43 24.8 E) on a 47 mm diameter, 0.22 μm pore sized PES filter (IT4IP, Belgium). Filters were flash frozen in liquid nitrogen. Mature river biofilms were collected after four weeks of natural colonization on glass slides (5 × 10 cm) on the *Gave de Pau* (43°24 14.2 N 0°37 30.4 W) water (Colas et al., 2023), then lyophilized prior to analysis. After homogenization, the different PMC strains biomasses, ground guts of the two fish species, and biofilms were split into six replicate subsamples by dividing the lyophilizates, three to be used for regular DNA extraction, and three for combined metabolite and DNA extraction. Filters from each of the three lakes were split in two halves, one for each extraction protocol.

### Metabolite extraction

Prior mechanical extraction (GLH850 OMNI; 25 000 r. min^-1^; 30s) was performed on all sample types except PMC strains. In all sample types, sonication (Sonics Vibra-Cell VCX 13; 60% amplitude; 30s) was then performed on weighted biomass or filter samples suspended in cold extraction solvent, consisting of 75-25% UHPLC methanol-water, 1 mL per 100 mg of tissue or per 10 mg of lyophilized culture, on ice (Colas et al., 2020). After centrifugation (10 min; 4 ° C; 15,300 g), remaining pellets were dried and used for subsequent DNA extraction.

### DNA extraction

DNA was extracted from the pellets obtained after metabolites extraction using the QIAGEN POWERLYZER POWERSOIL DNA extraction kit with a prior FastPrep 5G beat beater disruption (DNA Matrix; 5×30s; 8m.s^-1^) following manufacturer’s instructions (Foucault et al., 2022; Gallet et al., 2023). An extraction-blank control sample was also performed. Direct DNA extractions were also performed on the other three replicates of each sample type. In this case, DNA extraction was conducted using the same prior Bead-beating step, DNA extraction kit and protocol as above (but without prior extraction of metabolites).

### Sequencing and analysis of the V3-V4 region of prokaryotic 16S rRNA

The V3-V4 region of the 16S rRNA encoding gene was amplified using primers 341F (5’-CCTACGGGNGGCWGCAG -3’) and 806R (5_′_-GGACTACVSGGGTATCTAAT-3_′_) (Parada et al., 2016) and sequenced on an Illumina MiSeq 250×2 bp platform (GenoToul, Toulouse, France). Reads were deposited into the Sequence Read Archive (SRA) database (accession number PRJNA995525 (samples SRR25302593 -SRR25302641); Table S1).

Sequence analysis was performed using the QIIME2-2022.8 pipeline (Bolyen et al., 2019). Forward and reverse reads were trimmed at 230 and 225 bp, respectively. Amplicon Sequence Variants (ASVs) were obtained with the DADA2 plugin (default parameters) and affiliated with the SILVA 138-99 database. Diversity metrics were computed in RStudio (v4.1.3) with the R package phyloseq (v1.38.0) (McMurdie and Holmes, 2013). Statistical analyses were performed using R packages vegan (v2.6-4) (Oksanen et al., 2022) and pairwiseAdonis (v0.4) (Arbizu, 2022). All values are displayed as median ± standard deviation unless otherwise indicated.

## Results and discussion

### Non-significant impact of extraction method on diversity levels

Direct DNA extraction as well as sequential extraction of metabolites and DNA yielded similar PCR profiles with the primers used, and comparable numbers of reads in the Illumina sequencing (supplementary table 1). ASV richness, evenness and Shannon index obtained using both extraction procedures were not significantly different for a given sample type (Figure 1A-C, Kruskal-Wallis (KW) *p* > 0.05, supplementary tables 1 and 2). On the other hand, major differences were observed among sample types. Lowest richness was observed in non-axenic cyanobacterial strains, dominated by the *Microcystis aeruginosa* ASVs, which is congruent with recent reports indicating that these strains harbor a phycosphere that consists of a limited number of co-isolated or attached bacteria (Fu et al., 2020; Louati et al., 2015; Pascault et al., 2021; Seymour et al., 2017). Low richness was also observed in the medaka fish, congruent with recent reports (Duval et al., 2022) and the reported impact of domestication which usually leads to lower diversity compared to wild relatives (Alessandri et al., 2019; Hird, 2017). This is indirectly illustrated by the higher diversity level observed in the wild specimen of *Perca fluviatilis* collected from a lake compared to the medaka *Oryzias latipes* (Figure 1A-C; KW *p* < 0.05, Wilcoxon test (WT) *p* < 0.05; Supplementary tables 1 and 2). Highest diversity levels were observed in the Gave de Pau river’s biofilm sample that contain a quite diverse microbial community as previously observed for other biofilms (Romero et al., 2020), as biofilms are reported as hotspots of microbial diversity (Wu et al., 2019). Overall, recovered richness levels are thus congruent with the literature, and unaffected by the extraction method. This suggests that prior extraction of metabolites does not affect alpha diversity metrics in both low- and higher-diversity communities.

**Figure 1:**
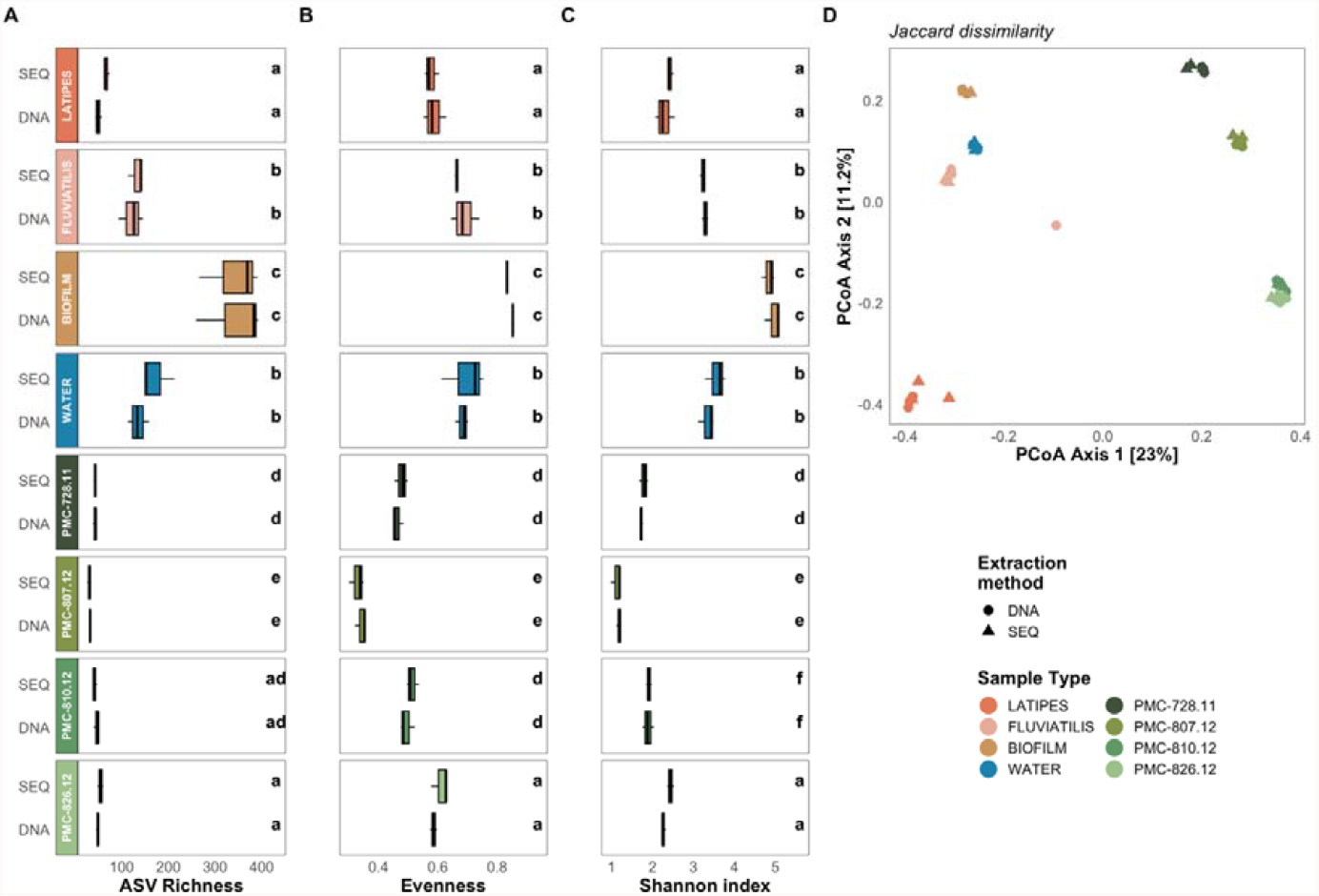
ASV richness (A), evenness (B) and Shannon index (C) for both extraction procedures, namely metabolites then DNA (sequential, SEQ) and classical DNA extraction (DNA) in the eight sample types (see methods). Median values and confidence intervals provided are based on three replicates per extraction type and sample type, with lowercase letters (a to e) on histograms indicating significance groups (Kruskall-Wallis test followed by a Wilcoxon post-hoc test). D: principal coordinates analysis of individual samples based on Jaccard distances, displaying explained variance percentage explained by the first two axes.

### Extraction method does not affect community composition and comparisons

Comparison based on Jaccard distances indicated that community compositions were highly similar among replicates from a given sample type and extraction method, as well as between the two extraction methods (DNA alone or sequential, Figure 1D; PERMANOVA p > 0.05; Supplementary table 2). Replicates from both extraction methods for each sample type formed tight clusters. The only exception was one of the three samples from *P. fluviatilis* for which DNA was extracted directly. It appeared outside of the cluster formed by the two others and the three on which sequential extraction was performed (Figure 1D). Clusters corresponding to each sample type were separated from each other except for clusters PMC-810.12 and PMC-826.12 which were overlapping. Because Jaccard distance is based on ASVs presence and absence, these results indicate that the two extraction methods yielded many identical ASVs, and thus a comparable image of the prokaryotic community. Bray-Curtis similarity plot, which also accounts for the abundance of these shared ASVs, yielded an overall similar picture (Figure 2A). One difference was that PMC-728.11 clustered with PMC-810.12 and PMC-826.12, suggesting a high abundance of the ASVs shared with the latter two communities.

**Figure 2:**
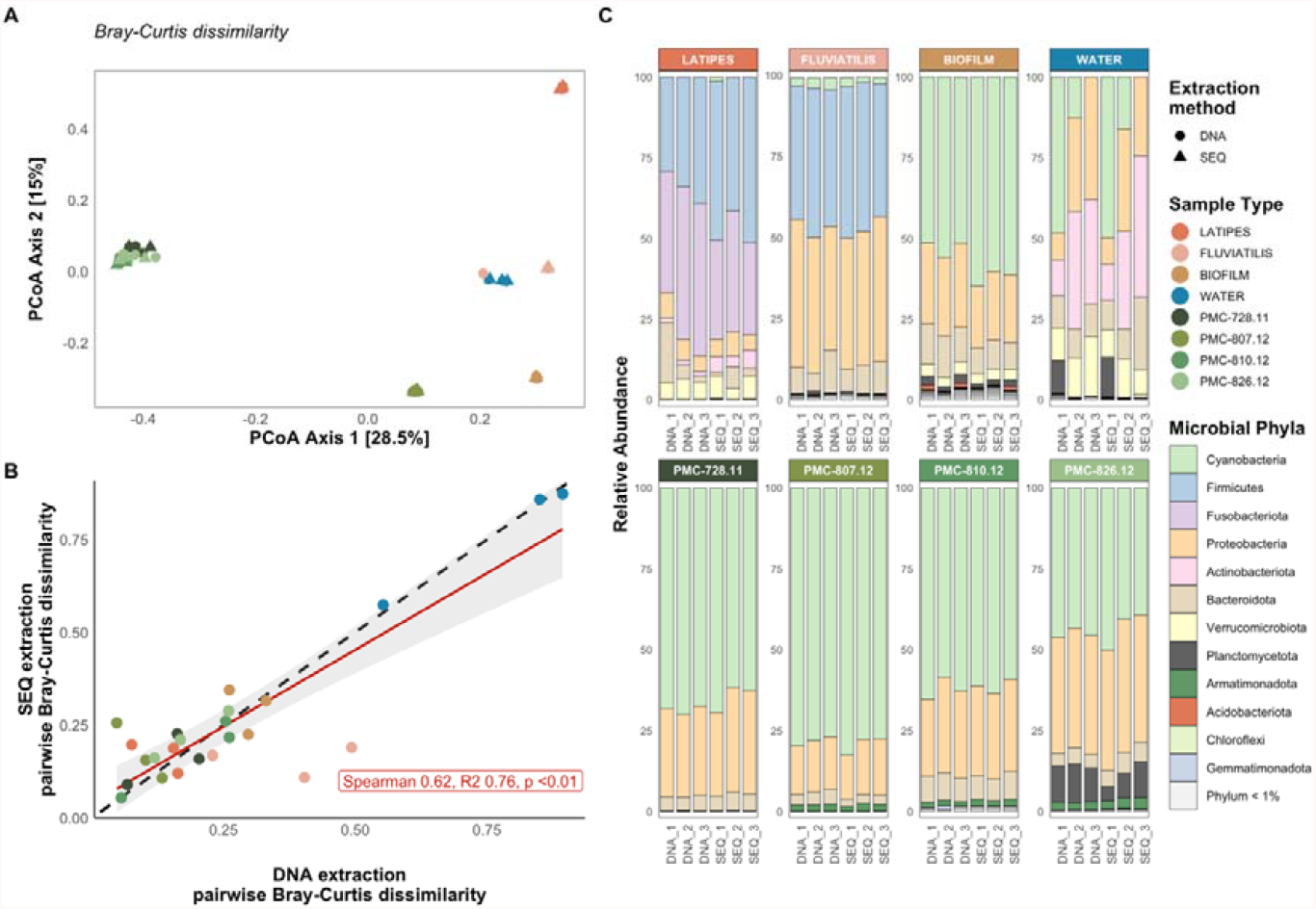
A: Principal coordinates analysis of individual samples based on Bray-Curtis distances, displaying explained variance percentage explained by the first two axes. B: Bray-Curtis pairwise dissimilarities between couples of samples within a given sample type computed from sequential (SEQ, y-axis) and DNA (DNA, x-axis) extractions. Red line corresponds to inferred linear relation, dashed black line corresponds to the 1:1 diagonal. C: prokaryotic community composition at the phylum level in each individual sample.

With the exception of the aforementioned *P. fluviatilis* sample, Bray-Curtis pairwise dissimilarities computed between couples of samples within a given sample type based on direct DNA extraction and sequential extractions were highly similar and correlated (Figure 2B; spearman 0.62, ANOVA R^2^ 0.76, *p* < 0.01; Supplementary table 2), suggesting that the extraction method had a very limited influence on inter-sample comparison metrics, and thus on our ability to properly compare community compositions. Compositions from individual replicates confirmed that they were very similar within a given sample type, regardless of the extraction method applied (Figure 2C). Various expected taxa typical of each sample type were identified, including for example abundant Firmicutes in fish samples (Llewellyn et al., 2014), Cyanobacteria, Proteobacteria, Bacteroidota and Verrucomicrobiota in peri-urban lakes (Louati et al., 2015), and the dominance of cyanobacteria in *Microcystis aeruginosa* strains as well as in the phototrophic biofilm. A similar picture was obtained at the genus level (Supplementary figure 1).

In conclusion, we found no evidence that prior extraction of metabolites had any influence on the result of prokaryotic community characterization using the V3-V4 region of the 16S rRNA-encoding gene and Illumina sequencing. This held true in a variety of sample types including low (strains) as well as high diversity communities (biofilm). These results indicate that metabolomic and 16S rRNA-based community characterization can be performed from a single sample, offering a unique opportunity to supplement the description of prokaryotic communities with a fingerprint of microbiome functioning. Single sample multi-omics are particularly relevant to microbiology because of the heterogeneity of most microbial habitats, the small scale at which we need to work, and potential cost- or biology-related limited sample availability. In the future, it will be important to confirm with additional sample types, 16S rRNA regions (Bukin et al., 2019), and other metabolite extraction mixes, but most importantly to test which additional types of omics could be performed from a single sample.

## Data availability

R scripts are available on Github upon publication (https://github.com/PierreFoucault/). All microbial communities’ samples sequencing reads were deposited into the NCBI Sequence Read Archive (SRA) database (accession number PRJNA995525 (samples SRR25302593 -SRR25302641); Supplementary Table 1).

## Acknowledgment

We thank Hydrosphere (Saint Ouen l’Aumône, France) for the sampling of wild perch, and Kandiah Santhirakumar who managed the medaka fish maintenance.

## Author’s contributions

BM and AG designed the sequential extraction protocol. SD and BM designed the study. CD, MG, AG, SC and PF performed the extractions and produced the DNA datasets. PF and SD analyzed the results. SD drafted the manuscript, all authors contributed and agreed with the latest version.

## Funding

The study was funded by Agence Nationale de la Recherche project COM2LIFE (ANR-20-CE32-0006). P.F. and M.G. are funded through the ANR project COM2LIFE, A.G. is funded through a Ph.D. grant from Ecole Doctorale 227 “Sciences de la Nature et de l’Homme”, MNHN.

## Ethics

Experimental procedures were carried out in accordance with European legislation on animal experimentation (European Union Directive 2010/63/EU) and were approved for ethical contentment by an independent ethical council (CEEA Cuvier n°68) and authorized by the French government under reference number APAFiS#19316-2019032913284201 v1. Fish were anesthetized in 0.1% tricaine methanesulfonate (MS-222; Sigma, St. Louis, MO) buffered with 0.1% NaHCO3 prior to sacrifice.

**Supplementary table 1:** sample IDs, SRA accession numbers, primer pairs used, raw and filtered (quality check, merging, chimera removal and taxonomy filtered) paired-end reads counts, metadata, Shannon, richness and evenness indices.

**Supplementary table 2:** Alpha and Beta diversity statistical test summaries.

**Supplementary figure 1:** prokaryotic community composition at the genus level in each individual sample. Only the 20 most abundant genera are displayed.

## Notes

### Competing Interest Statement

The authors have declared no competing interest.

## References

Alessandri, G., Milani, C., Mancabelli, L., Mangifesta, M., Lugli, G.A., Viappiani, A., Duranti, S., Turroni, F., Ossiprandi, M.C., van Sinderen, D., Ventura, M., 2019. The impact of human-facilitated selection on the gut microbiota of domesticated mammals. FEMS Microbiol. Ecol. 95. https://doi.org/10.1093/femsec/fiz121

Alivisatos, A.P., Blaser, M.J., Brodie, E.L., Chun, M., Dangl, J.L., Donohue, T.J., Dorrestein, P.C., Gilbert, J.A., Green, J.L., Jansson, J.K., Knight, R., Maxon, M.E., McFall-Ngai, M.J., Miller, J.F., Pollard, K.S., Ruby, E.G., Taha, S.A., Unified Microbiome Initiative Consortium, 2015. MICROBIOME. A unified initiative to harness Earth’s microbiomes. Science 350, 507–508. https://doi.org/10.1126/science.aac8480

Arbizu, P.M., 2022. pairwiseAdonis.

Bolyen, E., Rideout, J.R., Dillon, M.R., Bokulich, N.A., Abnet, C.C., Al-Ghalith, G.A., Alexander, H., Alm, E.J., Arumugam, M., Asnicar, F., Bai, Y., Bisanz, J.E., Bittinger, K., Brejnrod, A., Brislawn, C.J., Brown, C.T., Callahan, B.J., Caraballo-Rodríguez, A.M., Chase, J., Cope, E.K., Da Silva, R., Diener, C., Dorrestein, P.C., Douglas, G.M., Durall, D.M., Duvallet, C., Edwardson, C.F., Ernst, M., Estaki, M., Fouquier, J., Gauglitz, J.M., Gibbons, S.M., Gibson, D.L., Gonzalez, A., Gorlick, K., Guo, J., Hillmann, B., Holmes, S., Holste, H., Huttenhower, C., Huttley, G.A., Janssen, S., Jarmusch, A.K., Jiang, L., Kaehler, B.D., Kang, K.B., Keefe, C.R., Keim, P., Kelley, S.T., Knights, D., Koester, I., Kosciolek, T., Kreps, J., Langille, M.G.I., Lee, J., Ley, R., Liu, Y.-X., Loftfield, E., Lozupone, C., Maher, M., Marotz, C., Martin, B.D., McDonald, D., McIver, L.J., Melnik, A.V., Metcalf, J.L., Morgan, S.C., Morton, J.T., Naimey, A.T., Navas-Molina, J.A., Nothias, L.F., Orchanian, S.B., Pearson, T., Peoples, S.L., Petras, D., Preuss, M.L., Pruesse, E., Rasmussen, L.B., Rivers, A., Robeson, M.S., Rosenthal, P., Segata, N., Shaffer, M., Shiffer, A., Sinha, R., Song, S.J., Spear, J.R., Swafford, A.D., Thompson, L.R., Torres, P.J., Trinh, P., Tripathi, A., Turnbaugh, P.J., Ul-Hasan, S., van der Hooft, J.J.J., Vargas, F., Vázquez-Baeza, Y., Vogtmann, E., von Hippel, M., Walters, W., Wan, Y., Wang, M., Warren, J., Weber, K.C., Williamson, C.H.D., Willis, A.D., Xu, Z.Z., Zaneveld, J.R., Zhang, Y., Zhu, Q., Knight, R., Caporaso, J.G., 2019. Reproducible, interactive, scalable and extensible microbiome data science using QIIME 2. Nat. Biotechnol. 37, 852–857. https://doi.org/10.1038/s41587-019-0209-9

Bukin, Y.S., Galachyants, Y.P., Morozov, I.V., Bukin, S.V., Zakharenko, A.S., Zemskaya, T.I., 2019. The effect of 16S rRNA region choice on bacterial community metabarcoding results. Sci. Data 6, 190007. https://doi.org/10.1038/sdata.2019.7

Clerissi, C., Chaïb, S., Raviglione, D., Espiau, B., Bertrand, C., Meyer, J.-Y., 2023. Metabarcoding and Metabolomics Reveal the Effect of the Invasive Alien Tree Miconia calvescens DC. on Soil Diversity on the Tropical Island of Mo’orea (French Polynesia). Microorganisms 11, 832. https://doi.org/10.3390/microorganisms11040832

Colas, S., Duval, C., Marie, B., 2020. Toxicity, transfer and depuration of anatoxin-a (cyanobacterial neurotoxin) in medaka fish exposed by single-dose gavage. Aquat. Toxicol. 222, 105422. https://doi.org/10.1016/j.aquatox.2020.105422

Colas, S., Marie, B., Milhe-Poutingon, M., Lot, M.-C., Boullemant, A., Fortin, C., Faucheur, S.L., 2023. Meta-metabolomic Responses of River Biofilms to Cobalt Exposure and Use of Dose-response Model Trends as an Indicator of Effects. https://doi.org/10.1101/2023.06.19.545533

Dubilier, N., McFall-Ngai, M., Zhao, L., 2015. Microbiology: Create a global microbiome effort. Nature 526, 631–634. https://doi.org/10.1038/526631a

Duval, C., Marie, B., Foucault, P., Duperron, S., 2022. Establishment of the Bacterial Microbiota in a Lab-Reared Model Teleost Fish, the Medaka Oryzias latipes. Microorganisms 10, 2280. https://doi.org/10.3390/microorganisms10112280

Foucault, P., Gallet, A., Duval, C., Marie, B., Duperron, S., 2022. Gut microbiota and holobiont metabolome composition of the Medaka fish (Oryzias latipes) are affected by a short exposure to the cyanobacterium Microcystis aeruginosa. Aquat. Toxicol. 106329. https://doi.org/10.1016/j.aquatox.2022.106329

Fu, H., Uchimiya, M., Gore, J., Moran, M.A., 2020. Ecological drivers of bacterial community assembly in synthetic phycospheres. Proc. Natl. Acad. Sci. 117, 3656–3662. https://doi.org/10.1073/pnas.1917265117

Gallet, A., Halary, S., Duval, C., Huet, H., Duperron, S., Marie, B., 2023. Disruption of fish gut microbiota composition and holobiont’s metabolome during a simulated Microcystis aeruginosa (Cyanobacteria) bloom. Microbiome 11, 108. https://doi.org/10.1186/s40168-023-01558-2

Gertsman, I., Barshop, B.A., 2018. Promises and Pitfalls of Untargeted Metabolomics. J. Inherit. Metab. Dis. 41, 355–366. https://doi.org/10.1007/s10545-017-0130-7

Hamlaoui, S., Yéprémian, C., Duval, C., Marie, B., Djédiat, C., Piquet, B., Bernard, C., Duperron, S., 2022. The Culture Collection of Cyanobacteria and Microalgae at the French National Museum of Natural History: A Century Old But Still Alive and Kicking! Including in Memoriam: Professor Alain Couté. Cryptogam. Algol. 43, 41–83. https://doi.org/10.5252/cryptogamie-algologie2022v43a3

Hird, S.M., 2017. Evolutionary Biology Needs Wild Microbiomes. Front. Microbiol. 8. https://doi.org/10.3389/fmicb.2017.00725

Khemtong, C., Carpenter, N.R., Lumata, L.L., Merritt, M.E., Moreno, K.X., Kovacs, Z., Malloy, C.R., Sherry, A.D., 2015. Hyperpolarized 13C NMR detects rapid drug-induced changes in cardiac metabolism. Magn. Reson. Med. 74, 312–319. https://doi.org/10.1002/mrm.25419

Krassowski, M., Das, V., Sahu, S.K., Misra, B.B., 2020. State of the Field in Multi-Omics Research: From Computational Needs to Data Mining and Sharing. Front. Genet. 11.

Le Manach, S., Duval, C., Marie, A., Djediat, C., Catherine, A., Edery, M., Bernard, C., Marie, B., 2019. Global Metabolomic Characterizations of Microcystis spp. Highlights Clonal Diversity in Natural Bloom-Forming Populations and Expands Metabolite Structural Diversity. Front. Microbiol. 10. https://doi.org/10.3389/fmicb.2019.00791

Llewellyn, M.S., Boutin, S., Hoseinifar, S.H., Derome, N., 2014. Teleost microbiomes: the state of the art in their characterization, manipulation and importance in aquaculture and fisheries. Front. Microbiol. 5, 207. https://doi.org/10.3389/fmicb.2014.00207

Louati, I., Pascault, N., Debroas, D., Bernard, C., Humbert, J.-F., Leloup, J., 2015. Structural Diversity of Bacterial Communities Associated with Bloom-Forming Freshwater Cyanobacteria Differs According to the Cyanobacterial Genus. PloS One 10, e0140614. https://doi.org/10.1371/journal.pone.0140614

Louca, S., Polz, M.F., Mazel, F., Albright, M.B.N., Huber, J.A., O’Connor, M.I., Ackermann, M., Hahn, A.S., Srivastava, D.S., Crowe, S.A., Doebeli, M., Parfrey, L.W., 2018. Function and functional redundancy in microbial systems. Nat. Ecol. Evol. 2, 936–943. https://doi.org/10.1038/s41559-018-0519-1

Madaj, Z.B., Dahabieh, M.S., Kamalumpundi, V., Muhire, B., Pettinga, J., Siwicki, R.A., Ellis, A.E., Isaguirre, C., Escobar Galvis, M.L., DeCamp, L., Jones, R.G., Givan, S.A., Adams, M., Sheldon, R.D., 2023. Prior metabolite extraction fully preserves RNAseq quality and enables integrative multi-’omics analysis of the liver metabolic response to viral infection. RNA Biol. 20, 186–197. https://doi.org/10.1080/15476286.2023.2204586

McMurdie, P.J., Holmes, S., 2013. phyloseq: an R package for reproducible interactive analysis and graphics of microbiome census data. PloS One 8, e61217. https://doi.org/10.1371/journal.pone.0061217

Nothias, L.-F., Schmid, R., Garlet, A., Cameron, H., Leoty-Okombi, S., André-Frei, V., Fuchs, R., Dorrestein, P.C., Ternes, P., 2023. A multi-omics strategy for the study of microbial metabolism: application to the human skin’s microbiome. https://doi.org/10.1101/2023.03.26.532286

Nunan, N., 2017. The microbial habitat in soil: Scale, heterogeneity and functional consequences. J. Plant Nutr. Soil Sci. 180, 425–429. https://doi.org/10.1002/jpln.201700184

Oksanen, J., Simpson, G.L., Blanchet, F.G., Kindt, R., Legendre, P., Minchin, P.R., O’Hara, R.B., Solymos, P., Stevens, M.H.H., Szoecs, E., Wagner, H., Barbour, M., Bedward, M., Bolker, B., Borcard, D., Carvalho, G., Chirico, M., Caceres, M.D., Durand, S., Evangelista, H.B.A., FitzJohn, R., Friendly, M., Furneaux, B., Hannigan, G., Hill, M.O., Lahti, L., McGlinn, D., Ouellette, M.-H., Cunha, E.R., Smith, T., Stier, A., Braak, C.J.F.T., Weedon, J., 2022. vegan: Community Ecology Package.

Parada, A.E., Needham, D.M., Fuhrman, J.A., 2016. Every base matters: assessing small subunit rRNA primers for marine microbiomes with mock communities, time series and global field samples. Environ. Microbiol. 18, 1403–1414. https://doi.org/10.1111/1462-2920.13023

Pascault, N., Rué, O., Loux, V., Pédron, J., Martin, V., Tambosco, J., Bernard, C., Humbert, J.-F., Leloup, J., 2021. Insights into the cyanosphere: capturing the respective metabolisms of cyanobacteria and chemotrophic bacteria in natural conditions? Environ. Microbiol. Rep. 13. https://doi.org/10.1111/1758-2229.12944

Prasad Maharjan, R., Ferenci, T., 2003. Global metabolite analysis: the influence of extraction methodology on metabolome profiles of Escherichia coli. Anal. Biochem. 313, 145–154. https://doi.org/10.1016/S0003-2697(02)00536-5

Rauckhorst, A.J., Borcherding, N., Pape, D.J., Kraus, A.S., Scerbo, D.A., Taylor, E.B., 2022. Mouse tissue harvest-induced hypoxia rapidly alters the in vivo metabolome, between-genotype metabolite level differences, and 13C-tracing enrichments. Mol. Metab. 66, 101596. https://doi.org/10.1016/j.molmet.2022.101596

Romero, F., Acuña, V., Sabater, S., 2020. Multiple Stressors Determine Community Structure and Estimated Function of River Biofilm Bacteria. Appl. Environ. Microbiol. 86, e00291–20. https://doi.org/10.1128/AEM.00291-20

Roume, H., Heintz-Buschart, A., Muller, E.E.L., Wilmes, P., 2013. Sequential isolation of metabolites, RNA, DNA, and proteins from the same unique sample. Methods Enzymol. 531, 219–236. https://doi.org/10.1016/B978-0-12-407863-5.00011-3

Seymour, J.R., Amin, S.A., Raina, J.-B., Stocker, R., 2017. Zooming in on the phycosphere: the ecological interface for phytoplankton–bacteria relationships. Nat. Microbiol. 2, 1–12. https://doi.org/10.1038/nmicrobiol.2017.65

Shaffer, J.P., Nothias, L.-F., Thompson, L.R., Sanders, J.G., Salido, R.A., Couvillion, S.P., Brejnrod, A.D., Lejzerowicz, F., Haiminen, N., Huang, S., Lutz, H.L., Zhu, Q., Martino, C., Morton, J.T., Karthikeyan, S., Nothias-Esposito, M., Dührkop, K., Böcker, S., Kim, H.W., Aksenov, A.A., Bittremieux, W., Minich, J.J., Marotz, C., Bryant, M.M., Sanders, K., Schwartz, T., Humphrey, G., Vásquez-Baeza, Y., Tripathi, A., Parida, L., Carrieri, A.P., Beck, K.L., Das, P., González, A., McDonald, D., Ladau, J., Karst, S.M., Albertsen, M., Ackermann, G., DeReus, J., Thomas, T., Petras, D., Shade, A., Stegen, J., Song, S.J., Metz, T.O., Swafford, A.D., Dorrestein, P.C., Jansson, J.K., Gilbert, J.A., Knight, R., 2022. Standardized multi-omics of Earth’s microbiomes reveals microbial and metabolite diversity. Nat. Microbiol. 7, 2128–2150. https://doi.org/10.1038/s41564-022-01266-x

Singh, A., Shannon, C.P., Gautier, B., Rohart, F., Vacher, M., Tebbutt, S.J., Lê Cao, K.-A., 2019. DIABLO: an integrative approach for identifying key molecular drivers from multi-omics assays. Bioinforma. Oxf. Engl. 35, 3055–3062. https://doi.org/10.1093/bioinformatics/bty1054

Sogin, E.M., Puskás, E., Dubilier, N., Liebeke, M., 2019. Marine Metabolomics: a Method for Nontargeted Measurement of Metabolites in Seawater by Gas Chromatography–Mass Spectrometry. mSystems 4, 10.1128/msystems.00638-19. https://doi.org/10.1128/msystems.00638-19

Tarazona, S., Arzalluz-Luque, A., Conesa, A., 2021. Undisclosed, unmet and neglected challenges in multi-omics studies. Nat. Comput. Sci. 1, 395–402. https://doi.org/10.1038/s43588-021-00086-z

Valledor, L., Escandón, M., Meijón, M., Nukarinen, E., Cañal, M.J., Weckwerth, W., 2014. A universal protocol for the combined isolation of metabolites, DNA, long RNAs, small RNAs, and proteins from plants and microorganisms. Plant J. 79, 173–180. https://doi.org/10.1111/tpj.12546

Wu, Y., Cai, P., Jing, X., Niu, X., Ji, D., Ashry, N.M., Gao, C., Huang, Q., 2019. Soil biofilm formation enhances microbial community diversity and metabolic activity. Environ. Int. 132, 105116. https://doi.org/10.1016/j.envint.2019.105116

Zhang, X., Zhu, X., Wang, C., Zhang, H., Cai, Z., 2016. Non-targeted and targeted metabolomics approaches to diagnosing lung cancer and predicting patient prognosis. Oncotarget 7, 63437–63448. https://doi.org/10.18632/oncotarget.11521

