## Supplementary figure 1 for "Multi-omics analyses from a single sample: Prior metabolite extraction does not alter the 16S rRNA-based characterization of prokaryotic community in a diversity of sample types"

**Supplementary Figure 1:** Prokaryotic community composition at the Genus level in each individual sample. Only the 20 most abundant genera among all samples are displayed.

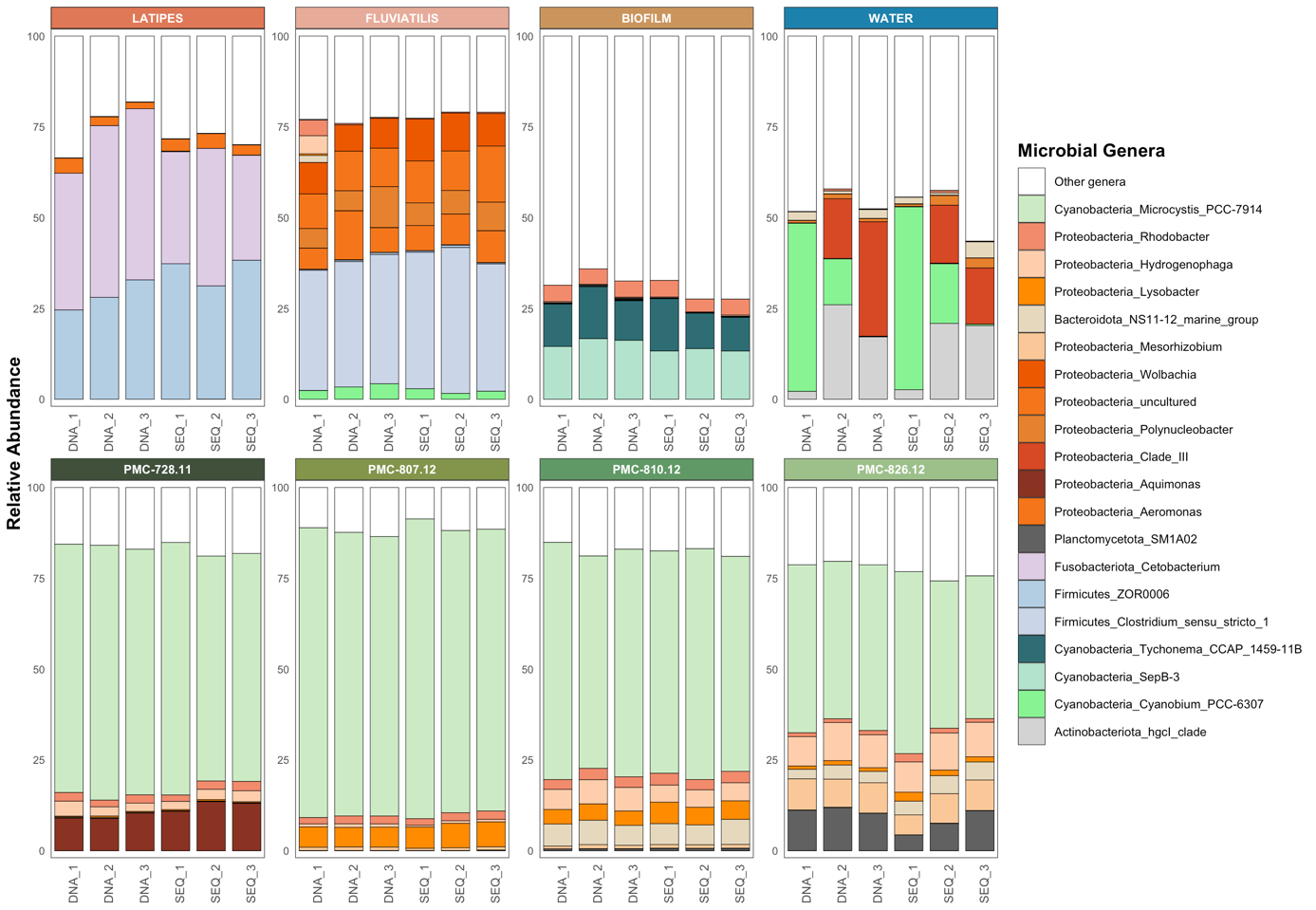
